# A receptor kinase complex refines cambium activity in *Arabidopsis*

**DOI:** 10.1101/2024.05.16.594489

**Authors:** Qing He, Jingyi Han, Wenbin Wei, Ehmke Pohl, Raymond Wightman, J Peter Etchells

## Abstract

In plant development, key receptor kinases are often active in disparate cell types, with each requiring vastly different signalling outputs. The ERECTA (ER) receptor kinase and its homologs ERL1 and ERL2 exemplify this pleiotropy. In *Arabidopsis*, they influence stomatal patterning, shoot meristem function, ovule morphogenesis, xylem fiber differentiation, and cell division in the vascular cambium^1–6^. Such diverse expression and functionality raises the question of how ER signalling can specify such distinct cell behaviours. One mechanism is via cell-type specific interactions with co-receptors, ligands, or other proteins that modulate signalling. However, little is known about ER interactors in the vascular cambium, a bifacial stem cell niche that generates phloem and xylem (**Figure 1A**). Combinatorial mutations between *ER, ERL1* and *ERL2* and receptor kinases of a second family, *PXY, PXL1*, and *PXL2*, show severe cambial defects^5,7^, but the mechanism underpinning these phenotypes is not known. Here we discovered that PXY proteins form protein complexes with ER and ERL2. PXY signalling can be manipulated by altering levels of its cognate ligand, TDIF. In genetic analysis, plant lines in which TDIF levels were altered had dramatic phenotypic changes that required the presence of ER or ERL2. Our results demonstrate that PXY signalling mediated cambium regulation depends on ER signalling and explains ER function in the cambium. Because the cambium produces xylem, which constitutes the wood in vascular plants, our findings position PXY-ER complexes at the centre of the accumulation of this versatile biomaterial and essential carbon sink.

## Results and discussion

### PXY and ER proteins interact at the plasma membrane

It has long been known that TDIF-PXY signalling is a key regulator of vascular development and that cambium initiation and maintenance are perturbed in its absence^8–10^, exemplified by reductions in cambial cell division in *pxy pxl1 pxl2* (*pxy-*family; *px*f) triple mutants^7,9^. These phenotypes are significantly enhanced by loss-of-function mutations in members of the ER family, *ER, ERL1*, and *ERL2* (*ER*-family; *ER*f)^3,5,7^. A high throughput Extracellular Interactome Assay (EICA) dataset which identified putative extracellular interactions between Arabidopsis receptor kinases, included interactions between the LRR domains of ER and PXY, ER and PXL1 and PXL2 and ERL2, albeit at low confidence levels^11,12^. We thus hypothesised that genetic interactions between *PXY* and *ER* family members might be underpinned by protein complex formation between the two receptor kinase families.

We used Förster Resonance Energy Transfer (FRET), which relies on the proximity of fluorophores with FRET being detected if two fluorophore-tagged proteins interact. *35S:PXY-CFP* and *35S::ER-YFP* constructs were generated and co-infiltrated into *Nicotiana* leaves. Epidermal cells expressing fluorescently labelled PXY and ER were analysed for co-localisation of signal and FRET. A YFP signal was observed upon excitation of CFP (**Figure 1B-E**), which was and measured at 21.4% indicating PXY-ER proteins interact (**Figure 1F**). We then determined if PXY could also complex with ERL1 and ERL2, by generating *35S::ERL1-YFP* and *35S::ERL2-YFP* constructs and co-infiltrating them with *35S:PXY-CFP*. A FRET signal was observed between PXY-CFP and ERL2-YFP which was measured with an efficiency of 21.8%. By contrast, we found no evidence for interactions between PXY and ERL1 (**Figure 1F, S1**).

**Figure 1.**
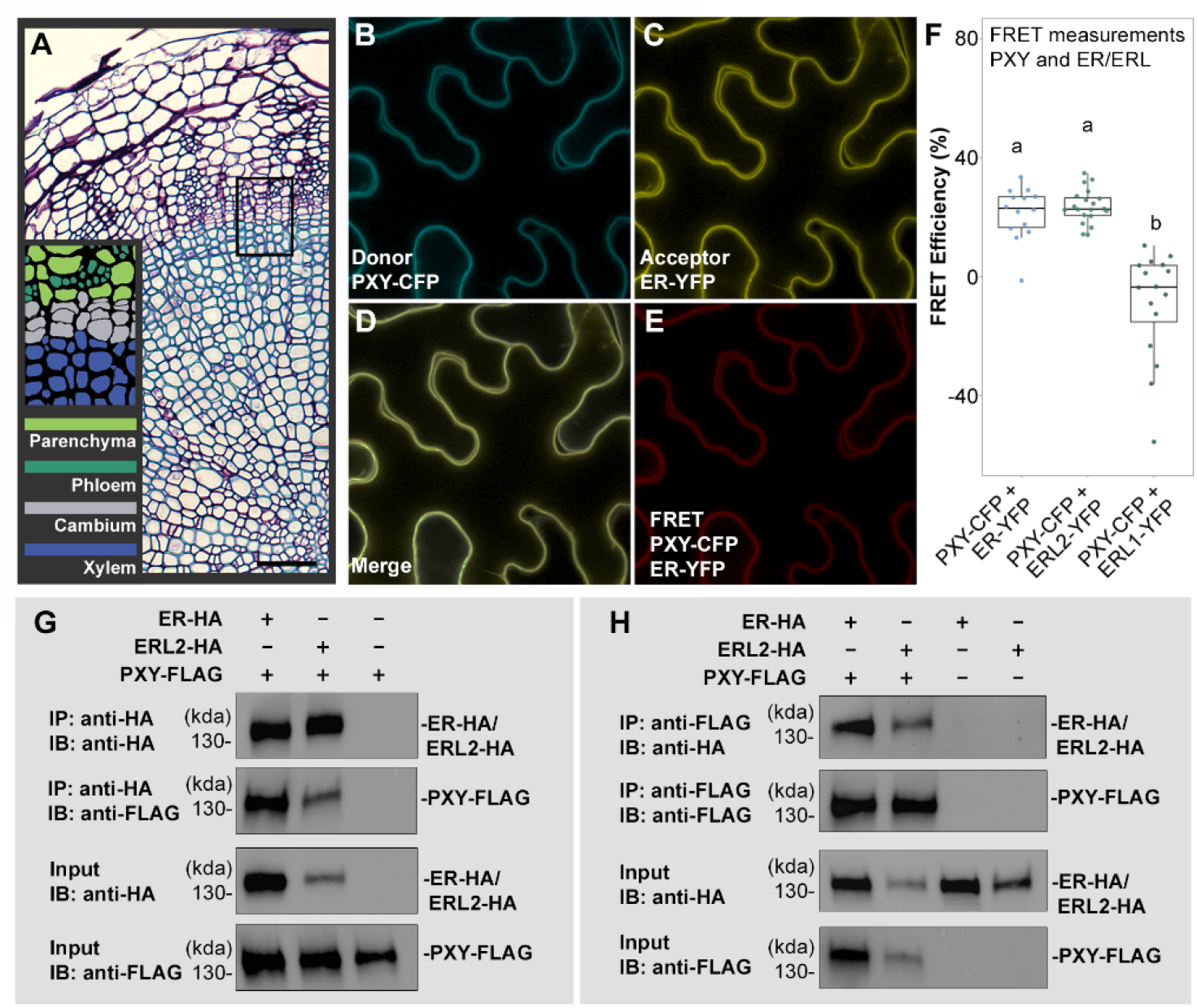
PXY forms heterodimers with ER and ERL2. **(A)** Transverse sections of a wild type hypocotyl showing the region of secondary growth. Black rectangle marks the area of cambium shown in the diagrammatic representation on the lower left. **(B-D)** Localisation of PXY-CFP (B), ER-YFP (C) and overlap (D) in *Nicotiana* epidermal cells. **(E)** FRET signal. **(F)** Graph showing FRET efficiencies of PXY-CFP and ER/ERL1/ERL2-YFP pairs. **(G-H)** Coimmunoprecipitation of PXY-FLAG with ER-HA or ERL2-HA and *vice versa* in *Arabidopsis* cells. Scales (A) are 50 μM.

Next, we used Co-immunoprecipitation (Co-IP) to further investigate the protein interactions observed between PXY and ER, and PXY and ERL2, again using transient expression in the *Nicotiana* system. Here, FLAG and HA epitope-tagged proteins were generated. A *35S:PXY-FLAG* translational fusion construct was co-infiltrated into *Nicotiana* leaves with *35S:ER-HA* or *35S:ERL2-HA*. PXY-FLAG was detected in pull downs using anti-HA beads, but not in *35S:PXY-FLAG* only controls (**Figure S2**). Similarly, when anti-FLAG beads were used ER-HA or ERL2-HA were pulled when *35S:PXY-FLAG* was co-infiltrated with *35S:ER-HA* or *35S:ERL2-HA*, but not in *35S:ER-HA* or *35S:ERL2-HA* only controls. As such, PXY-ER and PXY-ERL2 complexes can form in plants cells.

The EICA, FRET, and Co-IP experiments all supported the proposed protein interactions between PXY and ER or ERL2. However, all made use of either *in vitro* or heterologous systems. Consequently, Arabidopsis cell suspension cultures were used to determine whether PXY-ER or PXY-ERL2 heterodimers form in Arabidopsis cells. *35S:PXY-FLAG, 35S:ER-HA*, and *35S:ERL2-HA* constructs were generated and cells carrying either *35S:PXY-FLAG* and *35S:ER-HA*, or, *35S:PXY-FLAG* and *35S:ERL2-HA* were tested for interactions. ER-HA or ERL2-HA were detected in the presence of PXY-FLAG when protein extracts were co-incubated with anti-FLAG beads. Furthermore, PXY-FLAG was detected in the presence of ER-HA or ERL2-HA, when pulled down using anti-HA beads (**Figure 1G-H**). Thus, EICA, FRET, Co-IP in *Nicotiana* and Arabidopsis, collectively demon-strate that PXY heterodimerizes with ER and ERL2.

### Loss of ER attenuates PXY signalling

To understand the influence of PXY-ER/ERL2 complex formation on vascular development, we sought to perturb ER and ERL2 signalling in a sensitized PXY signalling background. TDIF, the PXY ligand, constitutes a dodecapeptide derived from *CLE41* and *CLE44. CLE41* and *CLE44* are predominantly phloem-expressed thus a gradient of TDIF forms across the cambium with a peak in the phloem^8,10^. By contrast, *PXY* expression predominates in the cambium-xylem boundary^8,9,13,14^. Consequently, expressing *CLE41* from a xylem-specific promoter results in phenotypes associated with a large increase in signalling though PXY receptors^10^. The *IRX3* promoter drives strong expression in differentiating xylem^15^. In wild type hypocotyls, cambial cell divisions are present in a narrow ring of tissue between xylem and phloem. Predominantly periclinal in nature, these cell divisions give rise to long cell files that align with the hypocotyl radial axis. By contrast ectopic cambium with misaligned cell divisions are apparent throughout the hypocotyl in *IRX3:CLE41* where this misalignment of cell division leads to very short cell files (**Figure 2A, D, G**).

**Figure 2.**
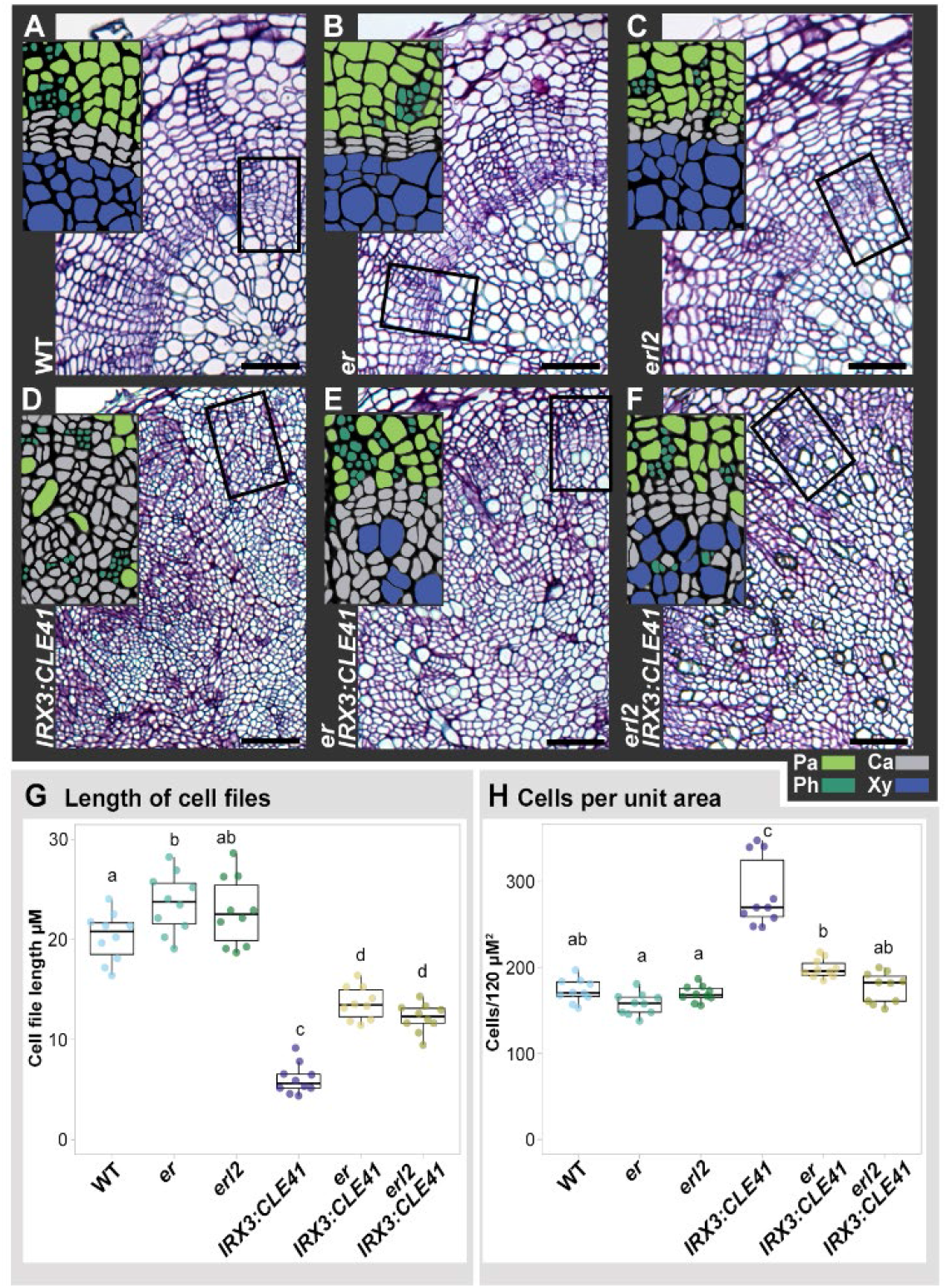
*er* and *erl2* suppress TDIF over-production phenotypes. **(A-F)** Transverse sections of hypocotyls. Thin sections show the extent of radial growth. Black rectangles mark area of cambium shown in diagrammatic representations on the left. Xylem and phloem are separated by cambium in wild type (A), *er* (B), and *erl2* (C). In *IRX3:CLE41* lines (D), very few xylem cells are apparent and the cambium is disordered. Organization of a cambial zone is restored in *er IRX3:CLE41* (E) and *erl2 IRX3:CLE41* (F) lines. **(G)** Boxplot showing length of cell files running parallel to the radial axis of the stem. **(H)** Boxplot showing cells per 120 μM^2^ in hypocotyl transverse sections. Letter above boxes mark significance groups (G and H; ANOVA + Tukey). Scales are 50 μM; Pa is parenchyma, Ca cambium, Ph phloem, and Xy xylem.

We tested whether ER and ERL2 were required for *IRX3:CLE41* phenotypes. *er IRX3:CLE41* and *erl2 IRX3:CLE41* lines were generated and compared to controls. In *er IRX3:CLE41* and *erl2 IRX3:CLE41* plants, aligned cell divisions in a defined cambial zone were present in a position comparable to the cambial zone of wild type plants (**Figure 2E-G**). Furthermore, average cell file length more than doubled in *er IRX3:CLE41* and *erl2 IRX3:CLE41* relative to *IRX3:CLE41* (**Figure 2E-G**), consistent with partial restoration of a spatially restricted cambium. Dividing cambium cells are typically smaller in size when compared to xylem vessels and parenchyma that occupy the centre of wild type hypocotyls. Consequently, an increase in cell size might be expected concomitant with restoration of spatially restricted cambium in *er IRX3:CLE41* and *erl2 IRX3:CLE41* lines, relative to *IRX3:CLE41* where the tissue was occupied predominantly with small cells reminiscent of actively dividing cambia. Indeed, cells in the centre of the hypocotyl of *er IRX3:CLE41* and *erl2 IRX3:CLE41* lines were larger than those observed in *IRX3:CLE41* lines. The number of cells per unit area was reduced by 31% in *er IRX3:CLE41*, and by 38% *erl2 IRX3:CLE41* when compared to *IRX3:CLE41* (**Figure 2H**). *er* and *erl2* thus partially suppressed *IRX3:CLE41* phenotypes confirming the importance of a functional ER signalling pathway to active TDIF-PXY signalling.

### ER is required for TDIF-PXY mediated repression of xylem differentiation

To better understand the processes governed by TDIF-PXY signalling to which the ER family contributes, transcriptomes of *IRX3:CLE41* and *er IRX3:CLE41* were obtained alongside wild type and *er* controls (**Supplemental dataset 1A**). Principal component analysis and clustering analysis were used to assess the level of similarity between the four genotypes tested. On the first principal component, which accounted for 76% of the variance between transcriptomes, *er IRX3:CLE41* was more similar to wild type and *er* than to *IRX3:CLE41* (**Figure 3A**). A similar result was observed using a Spearman correlation (**Figure S3A**), where again, *er IRX3:CLE41* clustered more closely with wild type and *er* than *IRX3:CLE41*, confirming earlier phenotypic analysis that *er* suppressed *IRX3:CLE41*. Consequently, differential gene expression was explored in further detail, with a focus on expression changes between *IRX3:CLE41* and *er IRX3:CLE41* (**Supplemental dataset 1B**). Gene Ontology (GO) analysis of biological function (**Supplemental dataset 1C**) supported our earlier observations which suggested that cell division was attenuated in *er IRX3:CLE41* relative the *IRX3:CLE41* (**Figure 2D-E, H**), as in a pairwise comparison between these two genotypes, GO categories enriched in *IRX3:CLE41* included several associated with cell division (**Figure S3**). PXY signalling has been shown to act as a positive regulator of transcriptional targets that promote cell division in the cambium. These include four cambium-expressed AINTEGUMENTA-like (AIL) transcription factors (*CAILs*; *ANT, AIL5*/*PLT5*, AIL6/*PLT3*, and *AIL7*/*PLT7*)^14^, two WUSCHEL RELATED HOMEOBOX transcription factors (*WOX4* and *WOX14*)^7,16^. WOX14 transcriptional targets include *TARGET OF MONOPTEROS 6* (*TMO6*) and *LATERAL ORGAN BOUNDARIES DOMAIN 4* (*LBD4*)^17^. Expression of all four *CAILS, WOX4, WOX14, TMO6*, and *LBD4* was lower in *er IRX3:CLE41* relative to *IRX3:CLE41* (**Figure S3C**). Thus, ER was required for the high expression of cambium cell-division promoting genes observed when TDIF levels were elevated in *IRX3:CLE41*.

**Figure 3.**
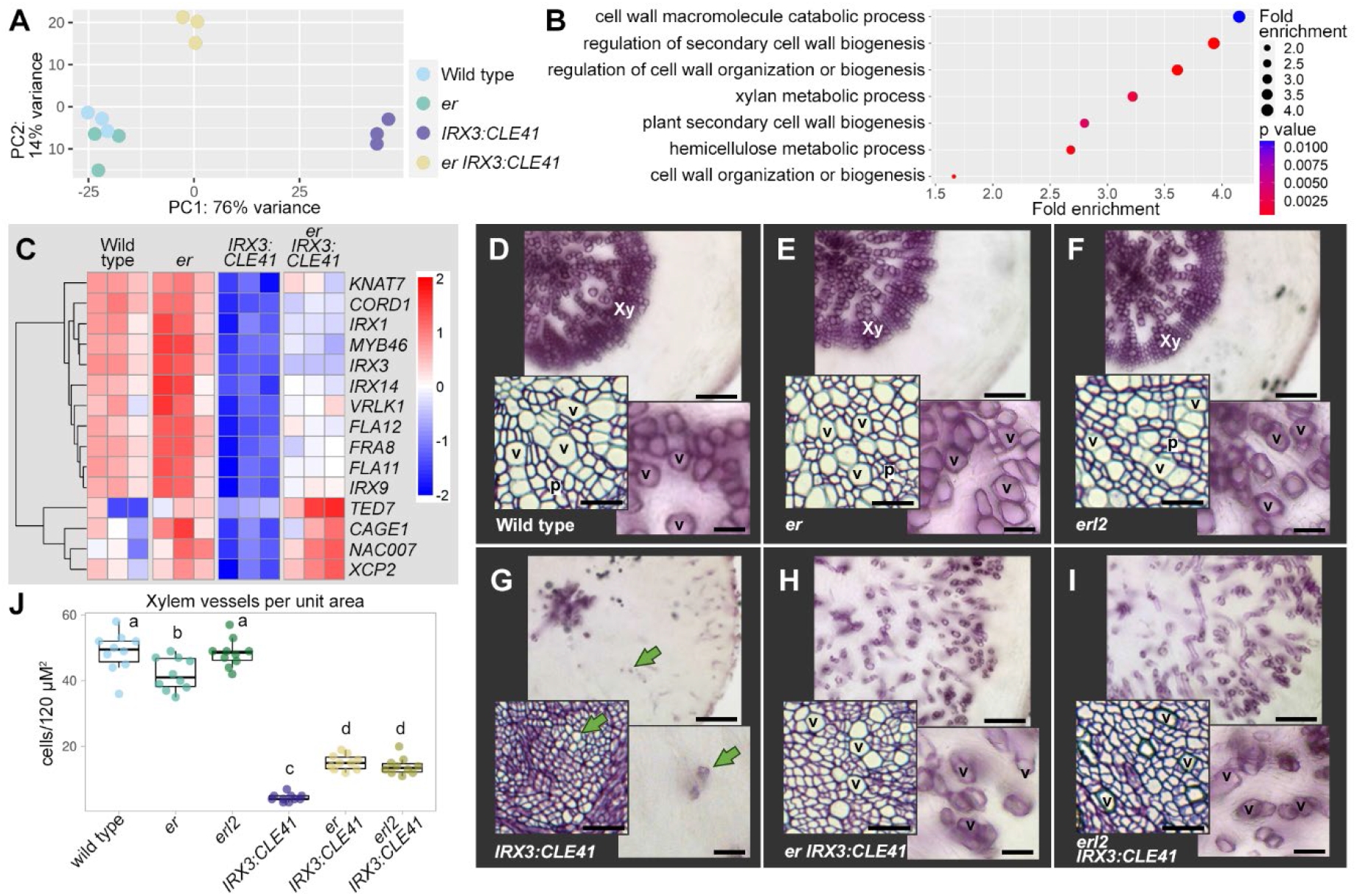
*er IRX3:CLE41* transcriptome. **(A)** Principal component analysis of wild type, *er, IRX3:CLE41* and *er IRX3:CLE41* transcriptomes demonstrates that of *er IRX3:CLE41* is more similar to that of wild type and *er* than *er IRX3:CLE41*. **(B)** Biological function ontogenies enriched in *er IRX3:CLE41* transcriptomes relative to *IRX3:CLE41*. (C) Heatmap showing expression of select xylem-enriched genes with differential expression in *er IRX3:CLE41* relative to *IRX3:CLE41*. (D-I) Transverse sections stained with phloroglucinol (upper and lower right panels) to visualise xylem differentiation with toluidine blue stained thin sections (lower left) for comparison. (J) Boxplot showing the number of xylem vessels per 120 μM^2^ in hypocotyl transverse sections. Letter above boxes mark significance groups (ANOVA + Tukey). Scales in D-I are 50 μM (upper panels) or 25 μM (lower panels); Xy is xylem, V marks vessels, green arrows point to vessel like cells in *IRX3:CLE41*.

Stem cell pools are generally maintained by controlled cell division, but also by exclusion of differentiation-promoting factors. Given that PXY promotes maintenance of the cambium stem cell pool, in part by excluding xylem differentiation from those cells^8,18^, it follows that the reduction in expression of genes associated with cell division and meristem maintenance in *er IRX3:CLE41* relative to *IRX3:CLE41* may be concomitant with increases in those involved in xylem formation. A hallmark of xylem vessel formation is deposition of a large secondary cell wall which is a composite of cellulose, hemicellulose and lignin, followed by programmed cell death^19^. GO categories enriched in *er IRX3:CLE41* relative to *IRX3:CLE41* included those associated with regulation and deposition of secondary cell walls (**Supplemental dataset 1D**). Furthermore, 23% of genes that demonstrated a higher expression in xylem cells relative to all other cell types in a high-resolution *Arabidopsis* root single cell transcriptome^20^ were up-regulated in *er IRX3:CLE41* relative to *IRX3:CLE41* representing a significant enrichment (p = 0.0031; **Supplemental dataset 1E**). Among the genes expressed at a higher level in *er IRX3:CLE41* relative to *IRX3:CLE41* were xylem differentiation regulatory genes *KNAT7, MYB46, NAC007*, and *VRLK1*; *CAGE1*, a cytoskeleton component that directs secondary cell wall deposition; enzymes that synthesise sugar polymers that constitute the cell wall, *IRX1, IRX3, IRX9, IRX14, FRA8, FLA11, FLA12, CORD1*; and *XCP* which promotes programmed cell death in xylem (**Figure 3C**). As such, ER is necessary to repress xylem differentiation in *IRX3:CLE41* lines.

During vegetative growth, Arabidopsis hypocotyl xylem is constituted of xylem vessels and parenchyma. Lignin is deposited in secondary cell walls such as those present on xylem vessels. To investigate xylem differentiation further, transverse sections were treated with phloroglucinol, which stains lignin subunits. Here, striking differences were observed between *IRX3:CLE41, er IRX3:CLE41*, and *erl2 IRX3:CLE41* (**Figure 3D-I**). Few cells were marked by phloroglucinol stain in *IRX3:CLE41*, but when either *ER* or *ERL2* was mutated, many were apparent, albeit fewer than was present in wild type or controls. Thin sections were taken to analyse the number of xylem vessels per unit area in *er IRX3:CLE41*, and *erl2 IRX3:CLE41* relative to controls (**Figure 3**). Consistent with phloroglucinol-stained tissue, these lines had more xylem elements than *IRX3:CLE41* but fewer than wild type, *er*, or *erl1* plants.

Collectively these results demonstrate that PXY signalling pathway is attenuated in the absence of ER and ERL2. PXY-ER heterodimers promote cambium cell division and exclude xylem differentiation in cambium, and as such these developmental outputs are attenuated in *er IRX3:CLE41*, and *erl2 IRX3:CLE41* plants presumably as PXY-ER or PXY-ERL2 multimers would be unable to form.

### Loss of the PXY-ER protein complexes results in ectopic xylem differentiation

Previously it had been shown that PXY can, in part, exclude xylem differentiation from the cambium^8,18^, and that combinatorial mutants within PXY and ER families result in a reduction in vascular tissue^5^. Having demonstrated that PXY forms complexes with members of the ER family (**Figure 1**) and that ER is necessary for repression of xylem differentiation when TDIF-PXY is constitutively active (**Figure 3**), we sought to bring together observations made in this paper with those in the literature^8,18^. Consequently, an analysis of xylem differentiation phenotypes when members of both PXY and ER families were absent was performed. *pxy pxl1 pxl2 er erl2* mutant transverse sections from 14-day old plants were analysed in comparison to wild type, *er erl2*, and *pxy pxl1 pxl2* controls (**Figure 4A**). Both vascular tissue area and the total number of vascular cells were reduced in *pxy pxl1 pxl2 er erl2* mutants (**Figure 4B-C**), but the number of xylem vessels remained unchanged across genotypes (**Figure 4D**). As such the proportion of xylem vessels as a total of the whole was significantly higher than the wild type, *er erl2*, or *pxy pxl1 pxl2* controls.

**Figure 4.**
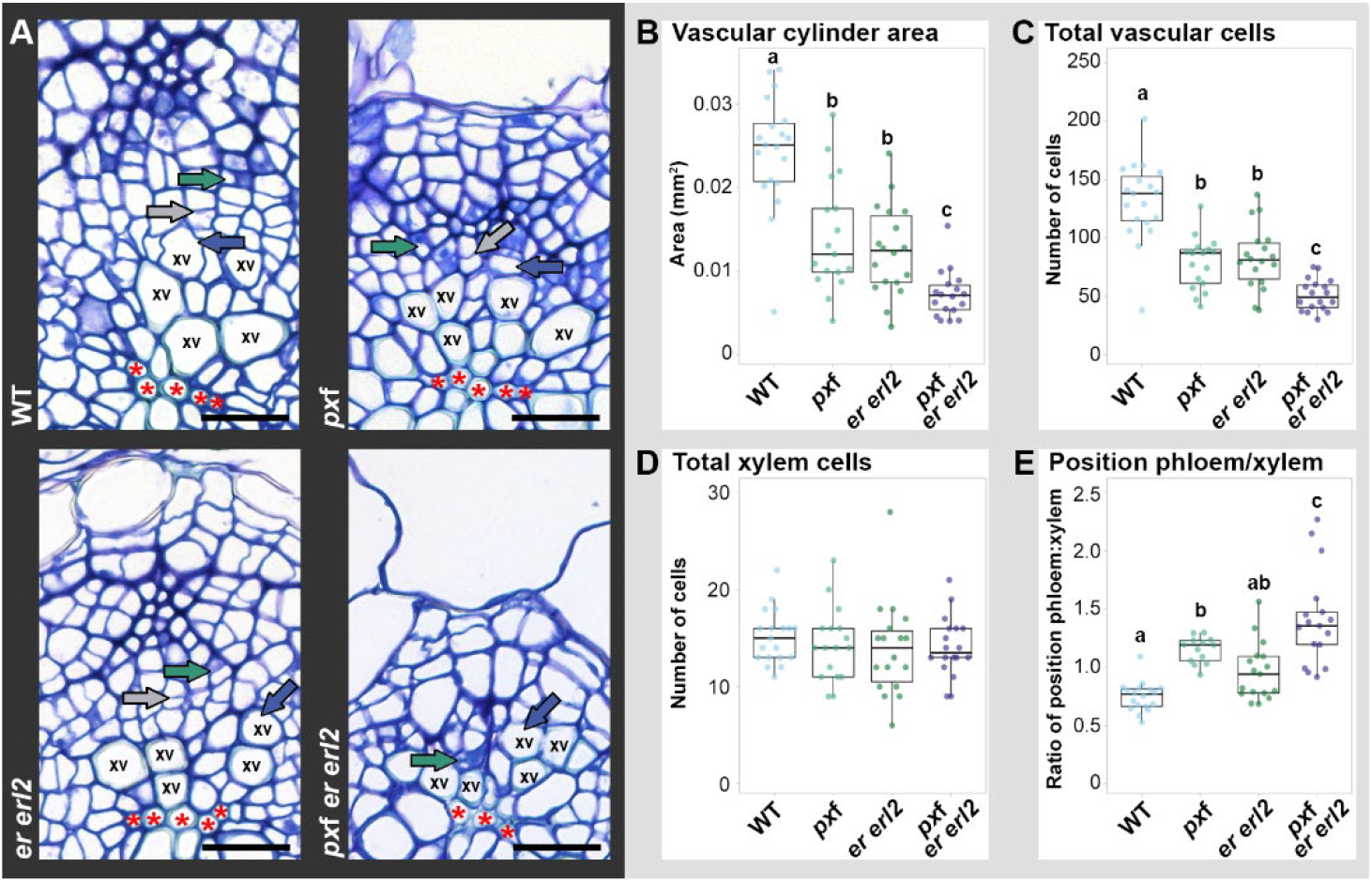
Loss of PXY and ER families results in failure to repress xylem differentiation. **(A)** Transverse sections of 14 day old hypocotyls of wild type, *pxy pxl1 pxl2* (*px*f), *er erl2*, and *px*f *er erl2* plants. **(B-D)** Boxplots showing area of vascular tissue in transverse sections (B), total number of vascular cells (C), and the number of xylem cells (D). **(E)** Boxplot showing ratios of the distance from central xylem axis to differentiating xylem: the distance from central xylem axis to differentiating phloem. Red asterisks mark the central xylem axis; xv is xylem vessel; green arrow denotes differentiation phloem; blue arrow is differentiation xylem; grey arrow marks the cambium. Letters above boxes in (B-E) mark significance groups (ANOVA + Tukey).

We then tested the position of xylem vessels relative to phloem. Since in wild type, xylem and phloem are derived from opposing sides of the cambium, with xylem forming towards the inside of the stem and phloem towards the outside (**Figure 1A**), the distance from the centre of the hypocotyl (**Figure 4A;** red asterisks) to differentiating xylem vessels (blue arrow in fig. 4A) is always smaller than the distance to differentiating phloem elements (green arrow in fig. 4A). As such the ratio of these distances is less than 1. We used this ratio as a proxy for xylem and phloem positioning, finding it to be higher in *pxy pxl1 pxl2 er erl2* mutants (ratio of 1.41) than in controls (**Figure 4E**). Consequently, the increased proportion of xylem vessels and their presence in ectopic positions (**Figure 4A**) are in support of PXY-ER complexes repressing xylem differentiation.

### Concluding remarks

The co-immunoprecipitation and FRET data shown here demonstrated that members of the PXY and ER families of receptor kinases form complexes at the plasma membrane (**Figure 1**). In genetic analysis, where PXY ligand was manipulated in the presence or absence of *er* or *erl2*, we have shown that PXY-ER heterodimers influence cell division (**Figure 2**) and repression of xylem differentiation (**Figure 3-4**), two key components required for cambium homeostasis. We propose that the interaction between PXY and ER defines ER outputs in the cambium. It remains an interesting question for future research as to the consequences of complex formation to signal transduction. ER has previously been shown to form complexes with receptor kinase SOBIR1 to control the rate of differentiation in xylem cells^21^, and with receptor-like protein TOO MANY MOUTHS to define epidermal patterning^22,23^, thus ER receptor heterodimerization with different tissue specific partners defines specific ER outputs within those tissues. In vascular tissue, there is abutment of tissues containing PXY-ER (cambium; **Figure 1**) and SOBIR1-ER (xylem) heterodimers^21^ likely collectively contribute to vascular tissue patterning and formation. Previously, we described loss of secondary growth in *pxy pxl1 pxl2 er erl2* lines, demonstrating a genetic interaction between members of the PXY and ER receptor families. Here we have shown that this genetic interaction underpins exclusion of xylem identity from the cambium via formation of PXY-ER receptor complexes.

## Acknowledgments

The authors are grateful to the Chinese Scholarship Council for funding a studentship to QH, and to BBSRC for funding grant number BB/V008129/1, to JH and JPE. This work was also supported by core microscopy facilities at Durham University and at SLCU. The Microscopy Core Facility at Sainsbury Laboratory, University of Cambridge, is supported by the Gatsby Charitable Foundation.

## Author contributions

Conceptualization, JPE, QH, EP; Methodology, RW, JPE, QH; Investigation, QH, WW, WW; Writing, JPE, QH; Visualization, QH, WW, JPE; Supervision, JPE, EP, RW; Funding Acquisition, QH, JPE, EP.

## Materials and methods

### Plant materials and Growth Condition

*Arabidopsis thaliana* (Columbia ecotype) plants were grown at 22 °C on a 16 h/8 h light/dark cycle. Seeds were surface-sterilized with absolute ethyl alcohol for 5 min and 75% ethanol for 15 min, then washed three-four times with sterilized distilled water under aseptic conditions. Surface-sterilized seeds were germinated on half-strength Murashige and Skoog (MS) medium in an incubator. After seven days, the seedlings were transferred into soil.

*er, erl2, pxy pxl1 pxl2 er erl2*, and *IRX3:CLE41* lines have been described previously^1–4^. *er IRX3:CLE41* and *erl2 IRX3:CLE41* were generated by crossing and selected in F2 and F3 populations on the basis of phenotype, PCR genotyping, and plant resistance.

*Nicotiana Benthamiana* seeds were germinated on soil and grown at 24°C16 h/8 h in a light/dark cycle.

### Binary vectors for FRET and Co-Immunoprecipitation

Primers (Table S1) were designed against *Arabidopsis thaliana PXY, PXL1, PXL2, ER, ERL1, ERL2* and used to amplify sequences from cDNA. Amplified cDNAs were subsequently fused with CFP/YFP/HA/FLAG tags via Infusion and cloned into pGWB502 using LR clonase II to generate constitutive expression vectors.

### FRET

Recombinant vectors *35S:PXY-CFP, 35S:PXL1-CFP, 35S:PXL2-CFP, 35S:ER-YFP, 35S:ERL2-YFP* and *35S:ERL2-YFP* were used for FRET. Vectors were transformed into tobacco leaf cells using Agrobacterium tumefaciens strain GV3101. 5ml of *Agrobacterium tumefaciens* strain GV3101 harbouring a binary vector was grown to an OD600 of 0.6, collected by centrifugation in 15ml tubes, washed twice and resuspended in 1ml infiltration buffer (1 M pH 5.7 MES 2.5 ml, 0.5 M D-Glucose 2.8 ml, 0.05 M Na^3^PO^4^ 12H^2^O 2 ml, 200 mM Acetosyringone 25 μl, with distilled water added to make a total volume of 50 ml) at room temperature. The final OD^600^ fell within the range of 0.01-0.1. cells from various constructs and P19^5^ were combined in a ratio of 1:1:1 (v/v/v). Transfected tobacco leaves were incubated in a greenhouse at 24 °C for a minimum of 48 h.

FRET imaging and quantification was carried out on a Zeiss LSM800 confocal microscope. FRET images were acquired in normal channel imaging mode using two GaAsP detectors and a Zeiss Objective Plan-Apochromat 63x/1.4 Oil DIC M27 (420782-9900-799) objective lens. A 458nm laser was used to excite the CFP/FRET detection and a 514 nm laser was used to excite the YFP. Detector 1 (Ch1) had an emission window set to 433-501 nm for CFP. Detector 2 (Ch2) had an emission window set to 500-566 nm for imaging the FRET/YFP. Calculation of FRET efficiencies used Lambda mode with the spectral detector set to a range of 409 to 607 nm and the 458 nm laser used as excitation source and a W Plan-Apochromat 40x/1.0 DIC M27 (421462-9900-799) objective lens. Scans were taking of CFP-YFP pairs and CFP only (that gives values of Donor contribution to the FRET channel). After acquiring a spectral scan in Zen software, regions of interest (ROIs, at the location of the plasma membrane of a transformed cell) were selected within the Spectral Unmixing tool. Values corresponding to the peaks of emission for CFP and YFP were used for subsequent FRET calculations. FRET efficiency was calculated using the formula: FRET Efficiency = Fa[minus the CFP contribution] / (Fd+Fa[minus the CFP contribution]).

### Co-immunoprecipitation

Co-immunoprecipitation was performed using either *Nicotiana* leaves infiltrated with *Agrobacterium* harbouring plant expression vectors, or *Arabidopsis* protoplasts. Protoplasts were isolated from root cell suspension cultures^6^ and then transfected with 10 μg plasmid DNA of the appropriate vectors and overnight dark incubated at room temperature in glucose-mannitol (GM) medium). Tissue was finely ground into a powder using mortar and pestles. Lysis binding buffer (50mM Tris-HCl, pH 7.6, 150mM NaCl, 1% NP-40, 0.1% SDS, 1mM PMSF, and Protease Inhibitor Cocktail - EDTA-Free) was added to the samples with a sample to buffer ratio between 1:1.5 and 1:2 (w/v). The resulting sample lysates were incubated on a shaker at 4 °C for 1 hour and centrifuged for 20 minutes at maximum speed and 4 °C.

Meanwhile, 30-40 μl of either Anti-DYKDDDDK or Anti-HA beads were placed on a magnetic rack, storage buffer removed, and washing buffer (consisting of 50mM Tris-HCl, pH 7.6, 150mM NaCl, 1% NP-40, 0.1% SDS, and 1mM PMSF) added. Beads to washing buffer ratio was maintained at 1:10 (v/v) though two further washes. 300-400 μl of sample lysate was introduced to the beads and incubated on a rocking table at 4 °C for 1-2 hours. Beads were washed 3-4 times as described earlier. Beads and 50-60 μl SDS-loading buffer were boiled at 94 °C for 5 minutes and then placed on a magnetic rack. Supernatants were subjected to western blots using standard methods.

### Plant anatomy

For thin sections stained with toluidine blue, plant material was fixed in FAA overnight and dehydrated through an ethanol series prior to embedding in JB4 (polysciences) according to the manufacturer’s instructions. Briefly, dehydrated samples were subjected to successive ethanol:JB4 infiltration solution 75%:25%, 50%:50%, 25%:75% solutions followed by 2 overnight incubations in 100% JB4 infiltration solution. Samples were dried on tissue, placed in moulds to which embedding solution was added. Blocks were covered with parafilm and hardened overnight. 4 μM sections were taken on a rotary microtome fitted with a glass knife, which were mounted on glass slides, stained with 0.025% aqueous toluidine blue and mounted with histomount.

Lignin was stained with phloroglucinol. Fresh sections embedded in 4% Agarose were taken using a vibratome which were incubated in phloroglucinol-HCl solution for 30 mins prior to mounting in glycerol for imaging.

### Gene expression

RNA was extracted from 5-week-old Arabidopsis hypocotyls in biological triplicate using ReliaPrep™ RNA Miniprep and subjected to RNA-seq on the Illumina platform (Novogene). 150 base paired end reads were mapped to the Arabidopsis TAIR10 genome (EnsemblePlants, release 58) sequence with corresponding gtf file to obtain reads per gene using STAR aligner (v 2.7.11a)^7^. Read count per gene was analysed using DESeq2(v 1.40.2)^8^ to get p values, adjusted p values, and log2 fold changes. Sample PCA (principal component analysis) plot was generated using the plotPCA function of DESeq2 after variance stabilization transformation. Sample correlation heatmap and gene expression heatmap were generated using pheatmap(v1.0.12)^9^. GO (gene ontology) analyses were performed using PANTHER^10^ and the results were plotted using ggplot2(v3.4.4)^11^. The raw sequencing dataset is available on the NCBI Gene Expression Omnibus (GEO) server under the accession number GSE263680.

### Accession numbers

PXY (At5g61480), PXL1 (At1g08590), PXL2 (At4g28650), ER (At2G26330), ERL1 (At5g62230), ERL2 (At5g07180), CLE41 (At3g24770), ANT (At4g37750), AIL5/PLT5 (At5g57390), AIL6/PLT3 (At5g10510), AIL7/PLT7 (At5g65510) WOX4 (At1g46480), WOX14 (At1g20700), TMO6 (At5g60200), LBD4 (At1g31320), BES1 (At1g19350), BZR1 (At1g75080), KNAT7 (At1g62990), CORD1 (At3g14170)), IRX1 (At4g18780), MYB46 (At5g12870), IRX3 (At5g17420), IRX14 (At4g36890), VRLK1 (At1g79620), FLA12 (At5g60490), FRA8 (At2g28110), FLA11 (At5g03170), IRX9 (At2g37090), TED7 (At5g48920), CAGE1 (At1g05170), NAC007 (At1g12260), XCP2 (At1g20850).

