## Supplemental figures for "A receptor kinase complex refines cambium activity in *Arabidopsis*"

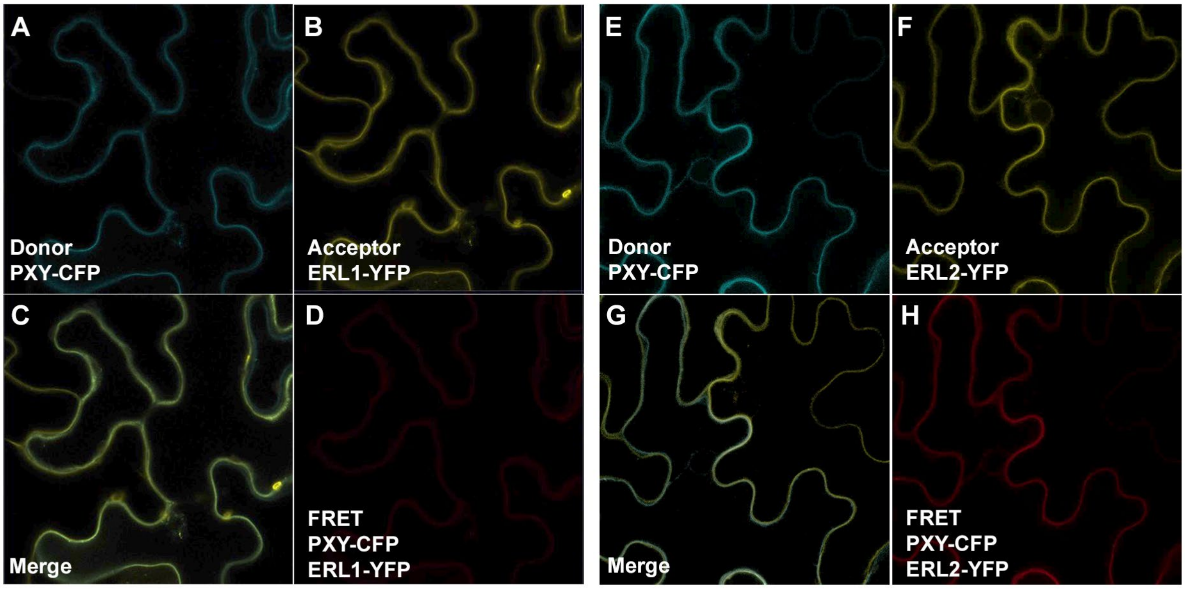


**Figure S1. FRET occurs between PXY-CFP and ERL2-YFP, but not between PXY-CFP and ERL1-YFP**
(**A-C**) Localisation of PXY-CFP (A) and ERL1-YFP (B) overlap (C) in *Nicotiana* epidermal cells. (**D**) No FRET signal was observed. (**E-G**) Localisation of PXY-CFP (E) and ERL2-YFP (F) overlap (G) in *Nicotiana* epidermal cells. (**H**) FRET signal.

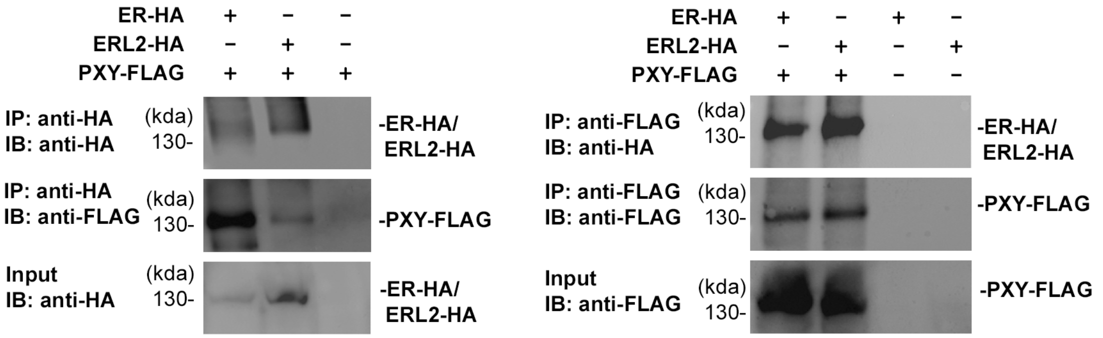


**Figure S2. PXY and ER/ERL2 heterodimerize in *Nicotiana* epidermis**
Coimmunoprecipitation of PXY-FLAG with ER-HA or ERL2-HA and *vice versa* in *Nicotiana* epidermis.

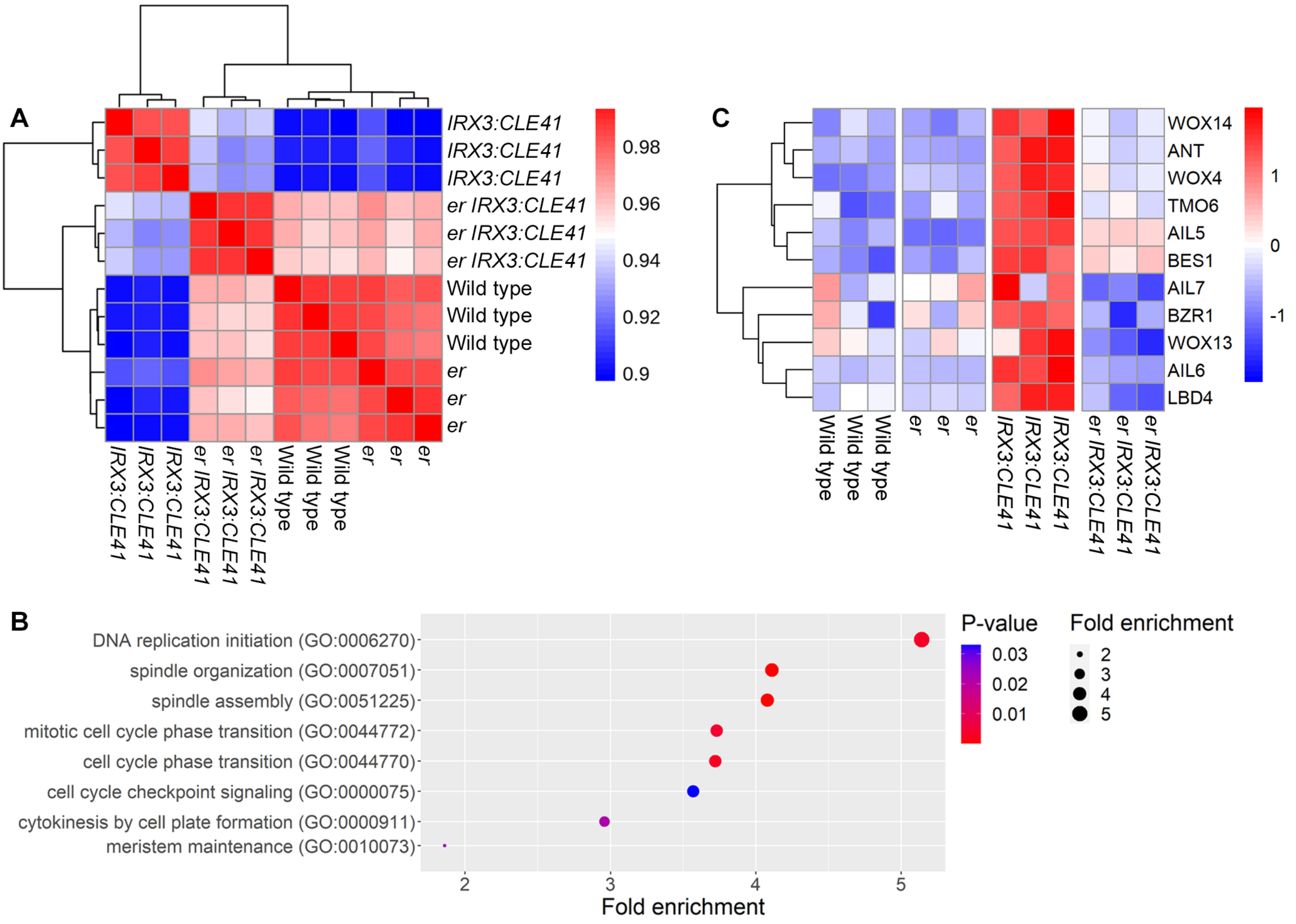


**Figure S3. Analysis of *er IRX3:CLE41* transcriptome**

(**A**) Cluster analysis of wild type, *er*, *IRX3:CLE41* and *er IRX3:CLE41* transcriptomes demonstrates that of *er IRX3:CLE41* is more similar to that of wild type and *er* than *er IRX3:CLE41*. (**B**) Biological function ontogenies associated with cell division enriched in *er IRX3:CLE41* transcriptomes relative to *IRX3:CLE41*. (**C**) Heatmap showing expression of transcriptional targets of TDIF-PXY signalling in *er IRX3:CLE41* relative to *IRX3:CLE41*.
